# Neural network dynamics underlying gamma synchronization deficits in schizophrenia

**DOI:** 10.1101/2020.10.07.330498

**Authors:** Daisuke Koshiyama, Makoto Miyakoshi, Yash B. Joshi, Juan L. Molina, Kumiko Tanaka-Koshiyama, Joyce Sprock, David L. Braff, Neal R. Swerdlow, Gregory A. Light

**Author notes:** **Corresponding Author:**, Gregory A. Light, Ph.D., Department of Psychiatry, University of California San Diego, Mail Code: 0804, 9500 Gilman Drive, La Jolla, CA, USA 92093-0804.

## Abstract

Gamma band (40-Hz) activity is associated with many sensory and cognitive functions, and is critical for cortico-cortical transmission and the integration of information across neural networks. The capacity to support gamma band activity can be indexed by the auditory steady-state response (ASSR); schizophrenia patients have selectively reduced synchrony to 40-Hz stimulation. While 40-Hz ASSR is a translatable electroencephalographic biomarker with emerging utility for therapeutic development for neuropsychiatric disorders, the spatiotemporal dynamics underlying the ASSR have not yet been characterized. In this study, a novel Granger causality analysis was applied to assess the propagation of gamma oscillations in response to 40-Hz steady-state stimulation across cortical sources in schizophrenia patients (n=426) and healthy comparison subjects (n=293). Results revealed distinct, hierarchically sequenced temporal and spatial response dynamics underlying gamma synchronization deficits in patients. During the response onset interval, patients exhibited abnormal connectivity of superior temporal and frontal gyri, followed by decreased information flow from superior temporal to middle cingulate gyrus. In the later (300–500 ms) interval of the ASSR response, patients showed significantly increased connectivity from superior temporal to middle frontal gyrus followed by broad failures to engage multiple prefrontal brain regions. In conclusion, these findings reveal the rapid disorganization of neural circuit functioning in response to simple gamma-frequency stimulation in schizophrenia patients. Deficits in the generation and maintenance of gamma-band oscillations in schizophrenia reflect a fundamental connectivity abnormality across a distributed network of temporo-frontal networks.

## Introduction

Information processing and psychophysiological measures have a long history in characterizing fundamental schizophrenia deficits underlying real word functioning.^1–4^ Recent advances in neuroscience have illuminated the time-linked mechanisms by which anatomically distinct brain regions communicate in order to integrate and coordinate perception, information processing, and cognition in mammals. In particular, the synchronous neural oscillations in the 30–50 Hz range that occur in the cortex, centered near 40 Hz and referred to as the “gamma band,” appear to reflect a fundamental central nervous system (CNS) resonance frequency critical for communication across multiple brain regions, allowing both basic and complex cognitive information processing to occur.^5–11^

Abnormalities in synchronous neural oscillations also help to explain cognitive deficits in complex brain illnesses like schizophrenia.^12–14^ Schizophrenia is characterized by information processing deficits in addition to a wide range of molecular, neurophysiological and neuroanatomic deficits. In schizophrenia, impairments in transient gamma synchrony measured by electroencephalography (EEG) are linked to not only basic and higher order cognitive deficits, but also clinical symptoms^15–18^; therefore, in schizophrenia, gamma band deficits are hypothesized to reflect the fundamental problems with “timing or sequencing component of mental activity” that lead to a heterogeneous constellation of clinical and cognitive deficits that characterize schizophrenia.^19^

The capacity to support the generation of synchronous neural activity can also be more directly interrogated by using paradigms that aim to “entrain” oscillations at particular frequencies. For example, studies of healthy subjects have revealed that when receiving periodic stimulation, neural networks behave like tuned oscillators, with EEG synchronizing to the frequency of stimulation.^20–22^ Schizophrenia patients have selectively reduced synchrony to auditory-based 40-Hz stimulation, but normal responses to other rates of stimulation.^3, 18, 23–36^ This gamma band auditory-steady response (ASSR) has received significant attention as a potential translatable biomarker advancing pro-cognitive treatment development in schizophrenia and other brain disorders, such as bipolar disorder, autism spectrum disorder and 22q11.2 deletion syndrome.^26, 35, 37–44^

Despite this interest, there is limited information in schizophrenia patients about how different brain regions coordinate, interact and produce ASSR deficits through the stimulus entrainment period.^45, 46^ Reports suggest the time course of ASSR may reflect distinct aspects of schizophrenia pathophysiology. Specifically, the 40-Hz ASSR is usually measured across a full 500 ms stimulation interval, with an early ~1–200 ms stimulus onset interval, and late ~300–500 ms maintenance interval^27, 28^: both first-episode as well as chronic schizophrenia patients have gamma-band ASSR deficits in early and later time intervals, but individuals at clinical high-risk for psychosis, showed deficits in only the later maintenance time interval.^25, 27, 28^

Augmenting magnetic resonance imaging (MRI)-based studies of *structural* connectivity, gamma-band ASSR can leverage the functional fine temporal resolution of EEG to assess *effective* connectivity, i.e. the causal interactions (i.e. directed information flows) among underlying brain regions that generate and maintain 40-Hz ASSR. This study aimed to use a novel, data-driven Granger causality analysis to deconstruct the time course and patterns of effective connectivity of cortical sources underlying the 40-Hz ASSR and its deficits in schizophrenia patients. This Granger causality approach enables an analysis of network dynamics with directed information flow using a correlation with time delay.

## Methods

### Subjects

Participants were 426 schizophrenia patients and 293 healthy comparison subjects (**Table 1**). Patients were recruited from community residential facilities and via clinician referral, and diagnosed using a modified version of the Structured Clinical Interview for DSM-IV-TR. Antipsychotic medications were prescribed for 388 schizophrenia patients. Healthy comparison subjects were recruited through internet advertisements. Exclusion criteria included an inability to understand the consent processes and/or provide consent or assent, not being a fluent English speaker, previous significant head injury with loss of consciousness, neurologic illness or severe systemic illness. Written informed consent was obtained from each subject. Audiometric testing was used to ensure that all participants could detect 1000-Hz tones at 40 dB. The Institutional Review Board of University of California San Diego approved all experimental procedures (071128, 071831, 170147). The current study was conducted in accordance with the Declaration of Helsinki.

**Table 1.** Demographic and clinical characteristics of subjects

|  | Healthy comparison subjects<br>(n = 293) |  | Schizophrenia patients<br>(n = 426) |  |
| --- | --- | --- | --- | --- |
|  | Mean | SD | Mean | SD |
| Gender (male/female) | 141/152 | – | 309/117 | – |
| Age (years) | 44.7 | 11.4 | 45.5 | 9.5 |
| Education (years) | 14.9 | 2.3 | 12.0 | 2.2 |
| Duration of illness (years)<sup>a</sup> | – | – | 23.7 | 10.1 |
| SAPS total score<sup>b</sup> | – | – | 8.9 | 4.1 |
| SANS total score<sup>b</sup> | – | – | 14.3 | 4.3 |
| GAF<sup>c</sup> | – | – | 40.9 | 6.6 |

Note: <sup>a</sup> Three subjects have no data of duration of illness; <sup>b</sup> Four subjects have no SAPS or SANS data; <sup>c</sup> One subject has no GAF data.
Abbreviations: SAPS, Scale for the Assessment of Positive Symptoms; SANS, Scale for the Assessment of Negative Symptoms.

### Stimuli and procedures

Auditory steady-state stimuli were presented to subjects by means of foam insert earphones (Model 3A; Aearo Company Auditory Systems, Indianapolis, Indiana). The stimuli were 1-millisecond, 93-dB clicks presented at 40 Hz in 500-millisecond trains. A block typically contained 200 trains of the clicks with 500-millisecond intervals. During the session, participants watched a silent cartoon video.

### Electroencephalography recording and preprocessing

EEG data were continuously digitized at a rate of 1000 Hz (nose reference, forehead ground) using a 40-channel Neuroscan system (Neuroscan Laboratories, El Paso, Texas). The electrode montage was based on standard positions in the International 10–5 electrode system^47^ fit to the MNI template head used in EEGLAB, including AFp10 and AFp9 as horizontal EOG channels, IEOG and SEOG above and below the left eye as vertical EOG channels, Fp1, Fp2, F7, F8, Fz, F3, F4, FC1, FC2, FC5, FC6, C3, Cz, C4, CP1, CP2, CP5, CP6, P7, P3, Pz, P4, P8, T7, T8, TP9, TP10, FT9, FT10, PO9, PO10, O1, O2, and Iz. Electrode-to-skin impedance mediated by conductive gel was brought below 4 kΩ. The system acquisition band pass was 0.5–100 Hz. Offline, EEG data were imported to EEGLAB 14.1.2^48^ running under Matlab 2017b (The MathWorks, Natick, MA). Data were high-pass filtered (FIR, Hamming window, cutoff frequency 0.5 Hz, transition bandwidth 0.5). EEGLAB plugin *clean_rawdata()* including Artifact Subspace Reconstruction was applied to reduce high-amplitude artifacts.^49-54^ The parameters used were: flat line removal, 10 s; electrode correlation, 0.7; ASR, 20; window rejection, 0.5. Mean channel rejection rate was 4.2 % (SD 2.3, range 0–15.8). Mean data rejection rate was 2.0% (SD 3.5, range 0–22.4). The rejected channels were interpolated using EEGLAB’s spline interpolation function. Data were re-referenced to average. Adaptive mixture ICA^55^ was applied to the preprocessed scalp recording data to obtain temporally maximally independent components (ICs). For scalp topography of each IC derived, equivalent current dipole was estimated using Fieldtrip functions.^56^ For scalp topographies more suitable for symmetrical bilateral dipoles, two symmetrical dipoles were estimated.^57^ To select brain ICs among all types of ICs, EEGLAB plugin *ICLabel()* was used.^58^ The inclusion criteria were 1) ‘brain’ label probability > 0.7 and 2) residual variance i.e., var((actual scalp topography) – (theoretical scalp projection from the fitted dipole))/var(actual scalp topography) < 0.15. Continuous EEG data were segmented into epochs that started at −250 ms and end at 750 ms relative to stimulus onset. Epoch rejection was performed for each participant using their median value of single-trial power spectral density (PSD) of remaining ICs. The single-trial PSD values were calculated and the median value across trials was subtracted. The residual errors from the median were z-scored. Any epoch with mean value > 2 between 15 to 35 Hz was rejected as contaminated by muscle potentials. The final processed data had a mean of 158.6 trials (SD 18.4).

### Effective connectivity analyses

To calculate grand-mean effective connectivity across ICs for each group, we applied EEGLAB plugin groupSIFT, which recently demonstrated successful application in other clinical EEG project.^59^ Using the same set of 9324 ICs in total preselected as representative brain ICs (mean 13.0 ICs per person, SD 4.4, range 4–28), time-frequency decomposed renormalized partial directed coherence (RPDC)^60^ was calculated across ICs (sliding window length 0.3 s, window step size 4 ms, logarithmically distributing 50 frequency bins from 2 to 55 Hz, baseline period −0.1 to 0 s). This generated a connectivity matrix with the dimension of IC × IC for each individual. The grand-average optimum model order was 11.6 (SD 1.6) i.e. delayed effective connectivity up to about 48 ms was utilized. The estimated equivalent dipole locations of the corresponding ICs were convolved with 3-D Gaussian kernel with 20 mm FWHM to obtain probabilistic dipole density. The dipole density inside the brain space is segmented into anatomical regions defined by automated anatomical labeling (AAL).^61^ The original AAL has 88 anatomical regions, but those basal and limbic regions that are unlikely to be scalp-measured EEG sources were integrated into two umbrella terms, upper basal and lower basal. The vagueness was intentional so that misleading use of limbic/basal label for EEG sources is avoided.

The original labels ‘upper basal’ and ‘lower basal’ were reported as ventral mid-cingulate, ‘mid-cingulate’ as dorsal mid-cingulate, and ‘insula’ as inferior frontal. This systematic bias toward depth in single-dipole fitting is due to the fact that ‘the large spatial extent of the many dipole layers that evidently generate EEG disqualifies them as single dipoles’.^62^ If this is the case, estimating shallower source regions along with radial direction as suggested here should provide a good heuristic correction. Individual IC × IC connectivity matrix is thus mapped to 76 × 76 custom anatomical region matrix, on which RPDC was also mapped as a weighting factor to modulate pairwise dipole density to calculate graph edges. For both groups, including a minimum of 80 % of unique subjects was set to be an inclusion criterion for each graph node to be analyzed in the next stage. Also, for the group comparison, 47/76 graph nodes showed overlap between the groups, which explained 79.3 % of total dipole density. For details of this solution, refer to our previous publication using this method.^59^ The analysis included 35–45 Hz as the frequency range of interest to be consistent with the frequency range in previous gamma-band ASSR studies.^27–30^ For the statistics of time-frequency decomposed RPDC within- and across-groups, weak family-wise error rate control was applied.^63, 64^

## Results

The connectivity matrix that represents the group-difference of gamma-band ASSR with the initial results *(p* < 0.05, corrected; two-tailed) is shown in **Figure 1**. A predefined *p*-value threshold of *p* < 0.0001 (two-tailed^65^) revealed 15 graph edges (**Figure 2**, **Supplementary Figure 1**. and **Supplementary Movie 1, 2)**. The matrix and movies of healthy comparison subjects (**Supplementary Figure 2** and **Supplementary Movie 3, 4)** and schizophrenia patients (**Supplementary Figure 3** and **Supplementary Movie 5, 6)** are shown in supplementary information.

**Figure 1.**
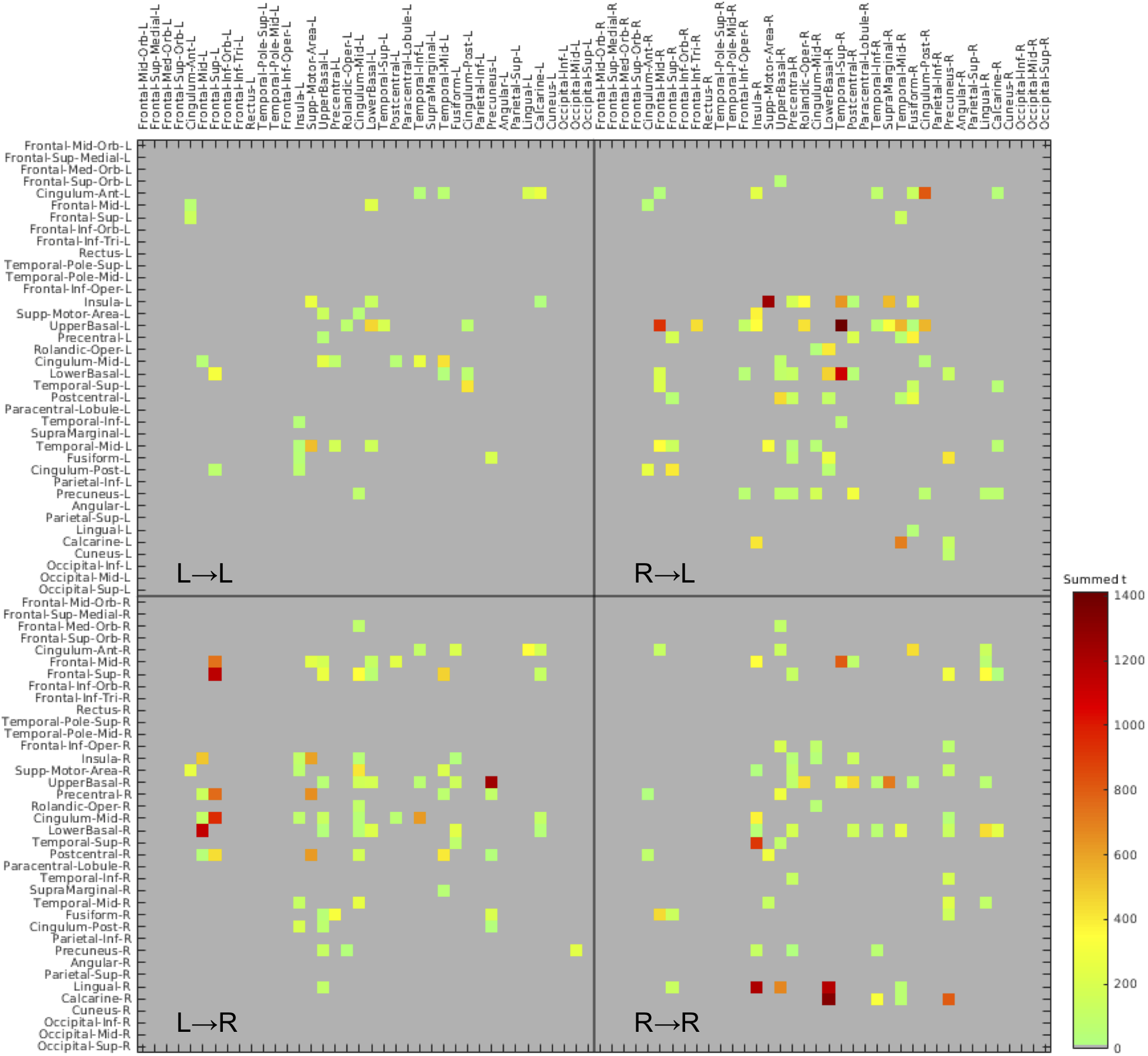
Connectivity matrix of 76 × 76 anatomical region of interests (ROIs) that represents the group-difference

**Figure 2.**
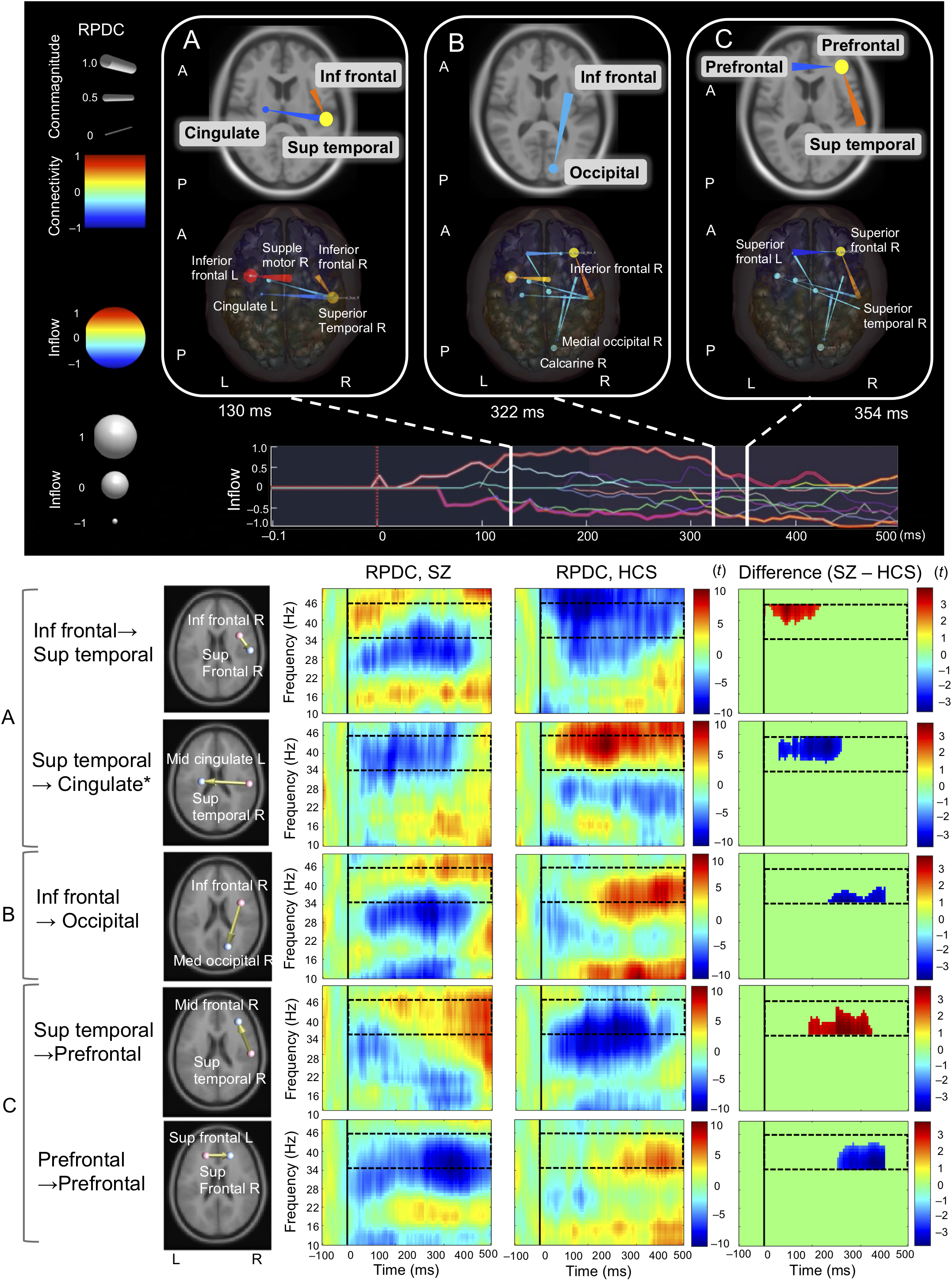
Difference of effective connectivity between schizophrenia and healthy comparison subject groups Note: Lower row of brain images indicates one frame of the effective connectivity movie at 130 ms (A), 322 ms (B) 354 ms (C) after the stimulus onset seen from an axial view (**Supplementary Movie 1** and **2**); upper row of brain images indicates main findings of connectivity in the flame; the panel shows the envelope of the significant edges between 35 and 45 Hz; lower part of figure shows the five major connectivity; solid black line at Latency = 0 is the stimulus onset; broken black line indicates region of interest between 35 and 45 Hz and between 0 and 500 ms; * the original label is ‘lower basal’ (refer to **Methods and Materials** section). Abbreviations: RPDC, renormalized partial directed coherence; Inf frontal; inferior frontal; Sup temporal; superior temporal; Supple motor area; supplementary motor area; SZ, schizophrenia; HCS, healthy comparison subject; A, anterior; P, posterior; L, left; R, right.

### Neural network dynamics

Based on the temporal dynamics, spatial interactions reflecting neural network dynamics underlying gamma synchronization deficits in patients are shown in **Figure 3A**. This network figure shows rank order latency of each information flow along with **Figure 3B**, revealing that the abnormal neural network centroid moves from an early local network centered at the right superior temporal to a late network centered at the left prefrontal in schizophrenia patients. Shortly after the response onset, a significant increase in connectivity was observed between temporal and frontal regions in schizophrenia patients (8 in **Figure 3B**), followed by decreased connectivity abnormalities within prefrontal regions (10, 12, 13 in **Figure 3B**), described below.

**Figure 3.**
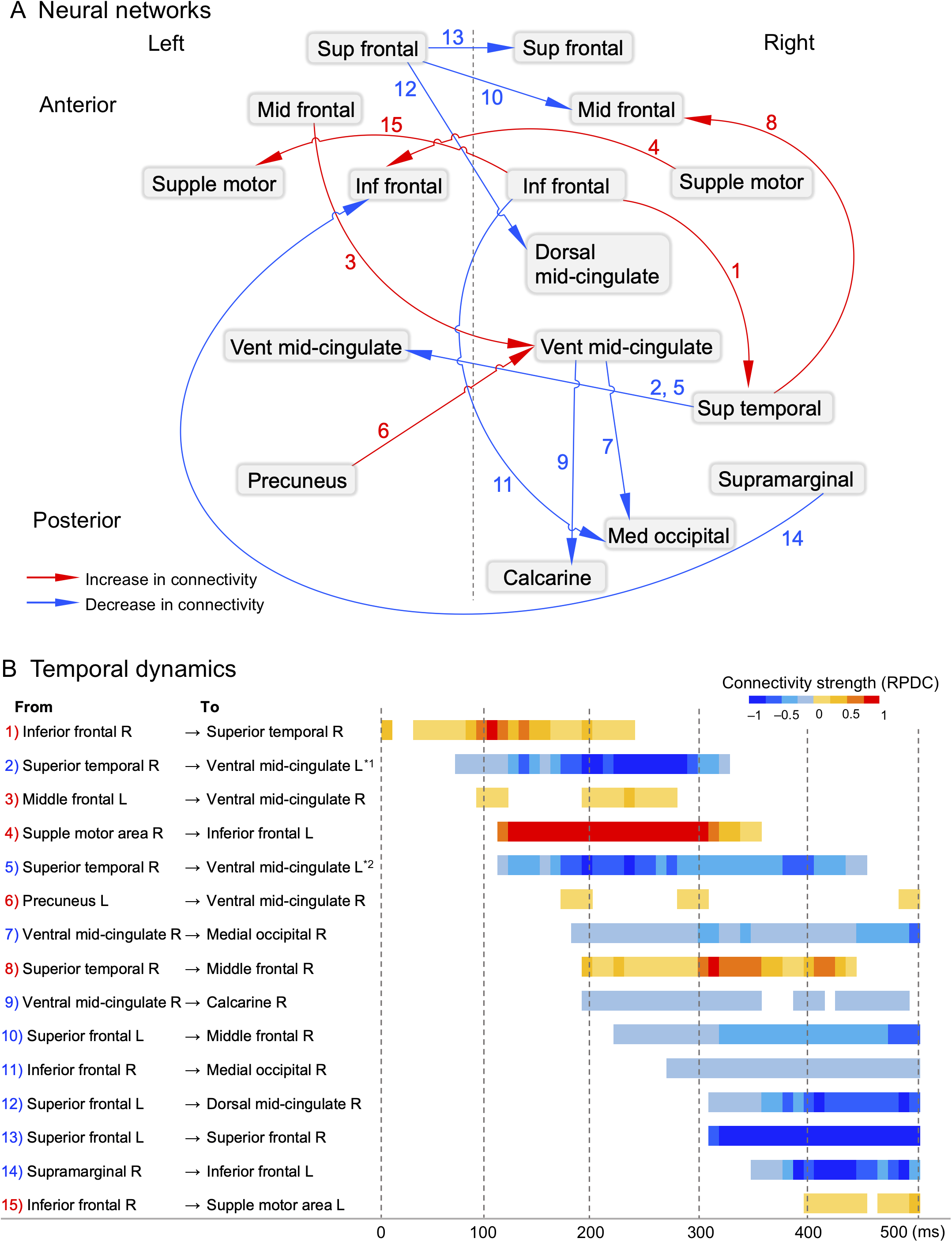
Neural networks (A) and temporal dynamics (B) underlying gamma oscillation deficits in schizophrenia patients Note: The numbers indicate rank order latency of each information flow; *^1^ the original label is ‘lower basal’; *^2^ the original label is ‘upper basal’ (refer to **Methods and Materials** section). Abbreviations: Sup frontal; superior frontal; Mid frontal; middle frontal; Supple motor, supplementary motor area; Inf frontal, inferior frontal; Dorsal mid-cingulate; dorsal middle cingulate; Vent mid-cingulate, ventral middle cingulate; Sup temporal; superior temporal; Med occipital, medial occipital; RPDC, renormalized partial directed coherence; L, left; R, right.

### Early ASSR time interval: network centered at superior temporal gyrus

Schizophrenia patients showed abnormal connectivity during the onset interval of ASSR (i.e., ~1–200 ms). Specifically, patients showed increased connectivity between right superior temporal and right inferior frontal gyri (**Figure 2A** and **Figure 3B**). Following response onset, information flow from the right superior temporal to the ventral middle-cingulate was decreased in patients compared to healthy comparison subjects (~100 ms to 300–400 ms). Patients also showed large increases in information flow from the right supplementary motor area to the left inferior frontal region compared to healthy comparison subjects.

### Middle time interval: information flow from inferior frontal to occipital regions

In the 200–300 ms response interval, connections from the right ventral middle cingulate to the right occipital regions, (i.e., right medial occipital and the right calcarine, right inferior frontal to the right medial occipital) were decreased in schizophrenia patients compared to healthy comparison subjects. These reductions continue into the later interval of ASSR (**Figure 2B** and **Figure 3B**).

### Later time interval: network centered at prefrontal regions

Schizophrenia related decreased connectivity during this later phase of ASSR (i.e., 300–500 ms) included the flows from the right parietal (supramarginal) to the left inferior frontal regions; this time interval was characterized by prominent and broad reductions in connectivity across prefrontal regions in schizophrenia patients (**Figure 2C** and **Figure 3B**).

## Discussion

Results of this study provide evidence of abnormal neural network dynamics underlying gamma synchronization deficits in schizophrenia patients compared to healthy comparison subjects. A novel Granger causality analysis was applied to EEG recordings obtained from large cohorts of schizophrenia patients and healthy comparison subjects ^66^. Gamma-band ASSR deficits in schizophrenia patients reflect abnormal neural network functioning centered at the superior temporal gyrus at the onset of the auditory steadystate response that then cascade “forward” and yield widespread abnormalities in the engagement of prefrontal brain regions in the later intervals of the response (**Figure 4**). Remarkably, before these decreases, abnormally increased information flows were evident in the local neural networks in patients (**Figure 3B**).

**Figure 4.**
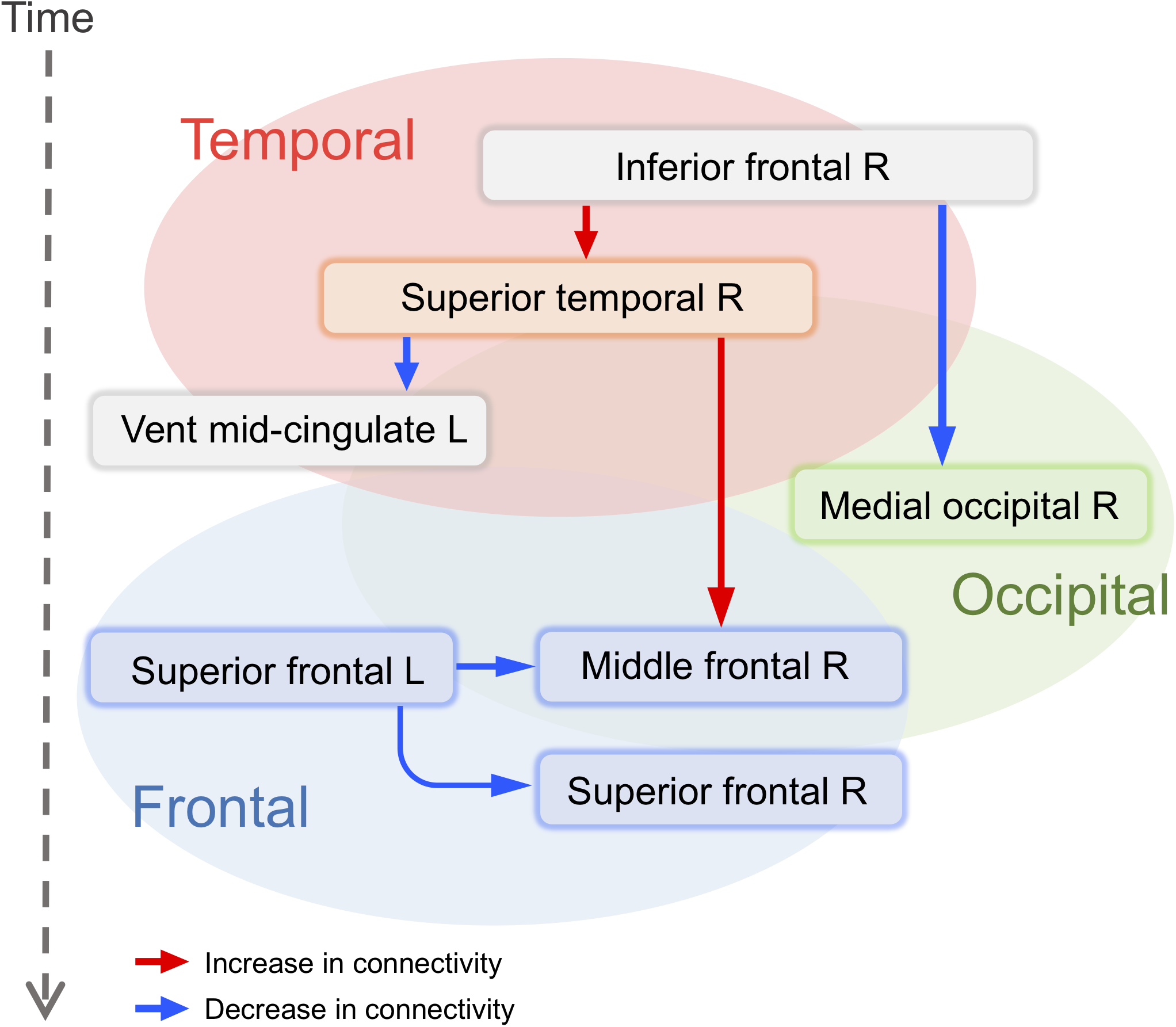
Abnormal neural network at the superior temporal gyrus in early phase and those networks at the prefrontal in late phase in schizophrenia patients

These results are strikingly consistent with a previous ECoG study by Tada et al.,^46^ which showed cortical surface levels of gamma-band ASSR in humans. They demonstrated prominent increases of gamma oscillations at the right primary auditory cortex (A1), right supramarginal gyrus and right supplementary motor area as well as moderate increases of activity at bilateral prefrontal gyri. While the Tada et al.^46^ findings were from patients undergoing neurosurgical intervention for treatment refractory epilepsy, they nonetheless suggest that these gamma-band locations are critical generators of ASSR in humans, and thus may be loci responsible for abnormalities in schizophrenia or other impaired neuropsychiatric patient populations.^46^ The current study, using novel and non-invasive EEG connectivity analyses may thus fill a gap between this study of patients with schizophrenia and the study of Tada et al. using ECoG and potentially may converge with findings from animal models.^46, 67^

The present study supports the increasing use of temporally distinct ASSR time intervals in studies of patients with psychosis. In this context, early and later time intervals of gamma oscillation appear to reflect dissociable response intervals.^25, 27, 28^ According to the prior studies, while first-episode schizophrenia patients showed gamma-band ASSR deficits both in the early and later intervals, individuals with clinical high-risks for psychosis showed reductions only in the later time interval. The local neural network centered at the superior temporal gyrus in the early onset time interval may be preserved in individuals with clinical high-risks for psychosis, consistent with the progressive volume reduction within the superior temporal gyrus observed after the onset of schizophrenia in first-episode schizophrenia patients.^68, 69^ The observed prefrontal gamma connectivity decrease during the later time interval in schizophrenia patients may reflect severe structural alterations in both gray and white matter.^70–73^

Support for the use of distinct time intervals also comes from an experimental medicine trial which revealed distinct pharmacodynamic results over the later maintenance, but not early onset phase of ASSR response in both healthy subjects and schizophrenia patients. Specifically, we reported that a single dose of the non-competitive N-methyl-D-aspartate (NMDA) receptor modulator, memantine, robustly increased (i.e., normalized) ASSR over later time intervals in schizophrenia patients.^37^

Results of this study should be considered in the context of some limitations. First, this is a cross-sectional cohort study of a heterogeneous sample of schizophrenia patients, the majority of whom were receiving complex medication regimens. As is the case for most large-scale studies of schizophrenia patients, the medication, psychosocial environments, and other important factors that could potentially influence brain function were not experimentally controlled. Second, the schizophrenia patients in this study had a well-established illness; results therefore may not generalize to at risk or early-illness psychosis patients. Third, 40 EEG channels were used for connectivity analyses. Future studies may fruitfully use higher density recordings with at least 64 channels,^74^ individual MRI data, and digitized scalp sensor locations rather than template head models and reliance on standardized electrode locations for potentially improved accuracy of source dynamics. Fourth, effective connectivity analyses are descriptive; they do not tell us about the functional roles of observed connectivity patterns or abnormalities; future studies will be needed to show their precise functional roles of the connectivity in neural substrates in patients with schizophrenia.

In conclusion, this study characterizes the disorganized propagation of gamma frequency oscillations in real-time in schizophrenia patients by applying a novel multivariate Granger causality analysis to large-scale scalp EEG datasets. Our findings provide evidence that relatively circumscribed abnormalities in the processing of simple 40-Hz auditory stimuli at the superior temporal gyrus in schizophrenia patients rapidly cascade forward to produce widespread failures to engage distributed prefrontal brain regions that are necessary to support gamma oscillations. Deficits in the generation and maintenance of stimulus-driven gamma-band oscillations reflect a fundamental connectivity abnormality across a distributed network of temporal and frontal regions. Clarification of the neural mechanisms underlying the networks detected in this study, in both future clinical and animal studies, will strengthen the utility of gamma-band ASSR as a translatable brain marker; this should further pave the way for clarifying the pathophysiology of neuropsychiatric and neurological diseases and the development of novel drug and behavioral therapeutic interventions.

## Acknowledgements

This study was supported by JSPS Overseas Research Fellowships (D. Koshiyama). Swartz Center for Computational Neuroscience is supported by generous gift of Swartz Foundation (New York; M. Miyakoshi). The funders had no role in the study design, data collection and analysis, publication decision, or manuscript preparation.

## Supplementary Information

### TABLE OF CONTENTS

**SUPPLEMENTARY FIGURES**

**Supplementary Figure 1** Increased and decreased connectivity in schizophrenia

**Supplementary Figure 2** Connectivity matrix in healthy comparison subjects

**Supplementary Figure 3** Connectivity matrix in schizophrenia patients

### SUPPLEMENTARY MOVIES (Only title and legends)

**Supplementary Movie 1** Difference of effective connectivity between schizophrenia and healthy comparison subject groups seen from an axial view

**Supplementary Movie 2** Difference of effective connectivity between schizophrenia and healthy comparison subject groups seen from a sagittal view

**Supplementary Movie 3** Effective connectivity in healthy comparison subject group seen from an axial view

**Supplementary Movie 4** Effective connectivity in healthy comparison subject group seen from a sagittal view

**Supplementary Movie 5** Effective connectivity in schizophrenia group seen from an axial view

**Supplementary Movie 6** Effective connectivity in schizophrenia group seen from a sagittal view

**Supplementary Figure 1.**
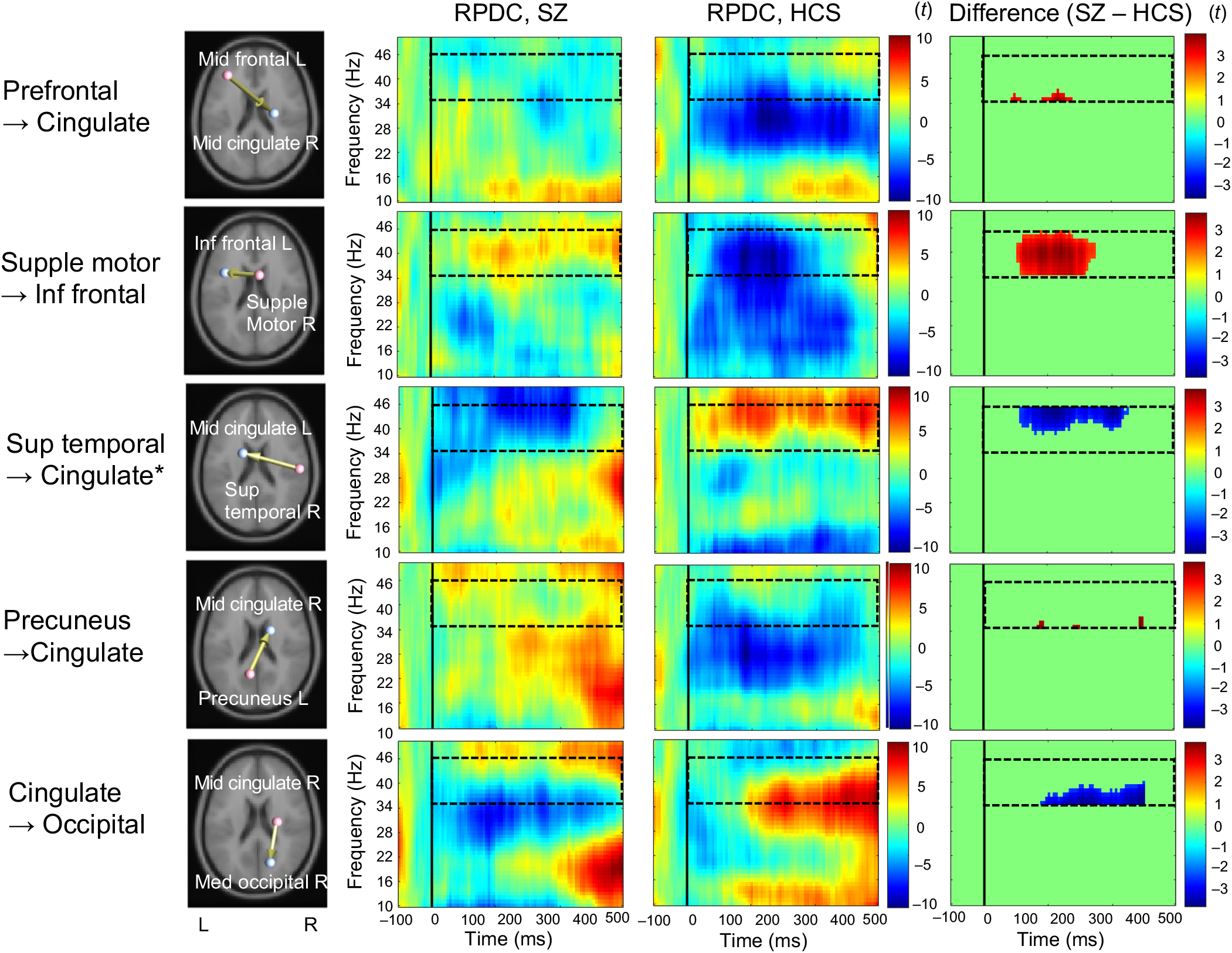
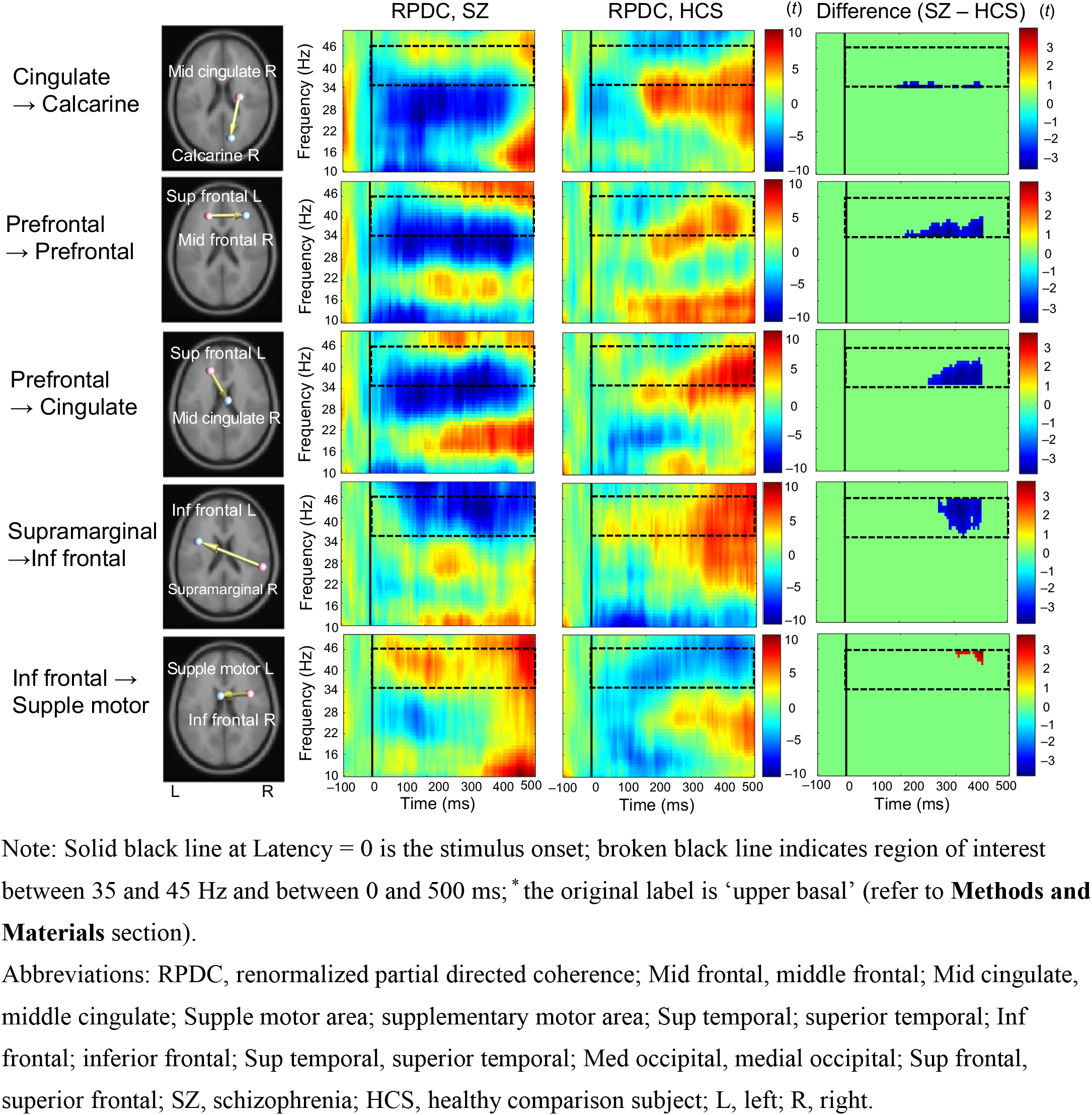
Increased and decreased connectivity in schizophrenia. Abbreviations: RPDC, renormalized partial directed coherence.

**Supplementary Figure 2.**
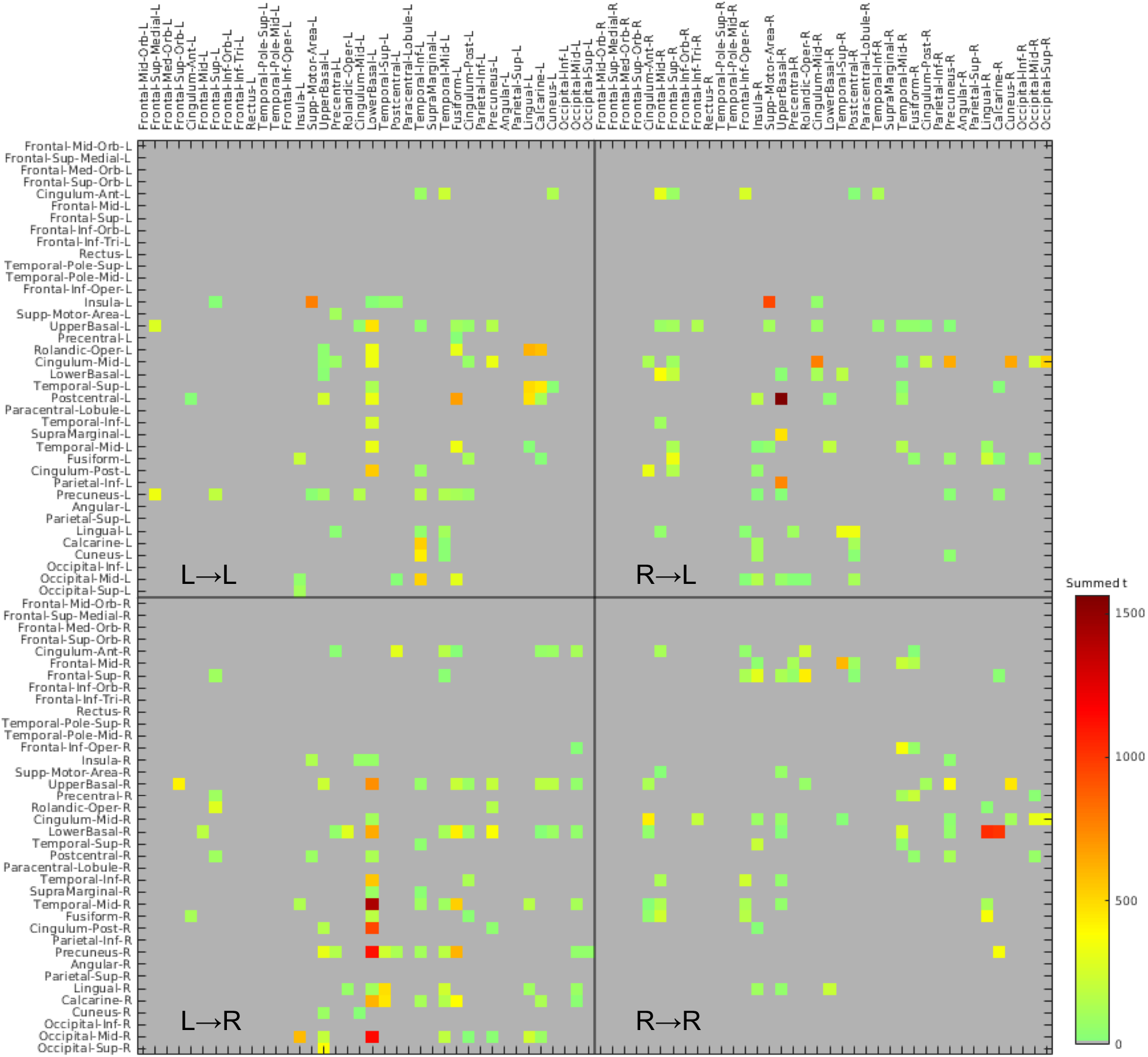
Connectivity matrix of 76 × 76 anatomical regions of interests in healthy comparison subjects

**Supplementary Figure 3.**
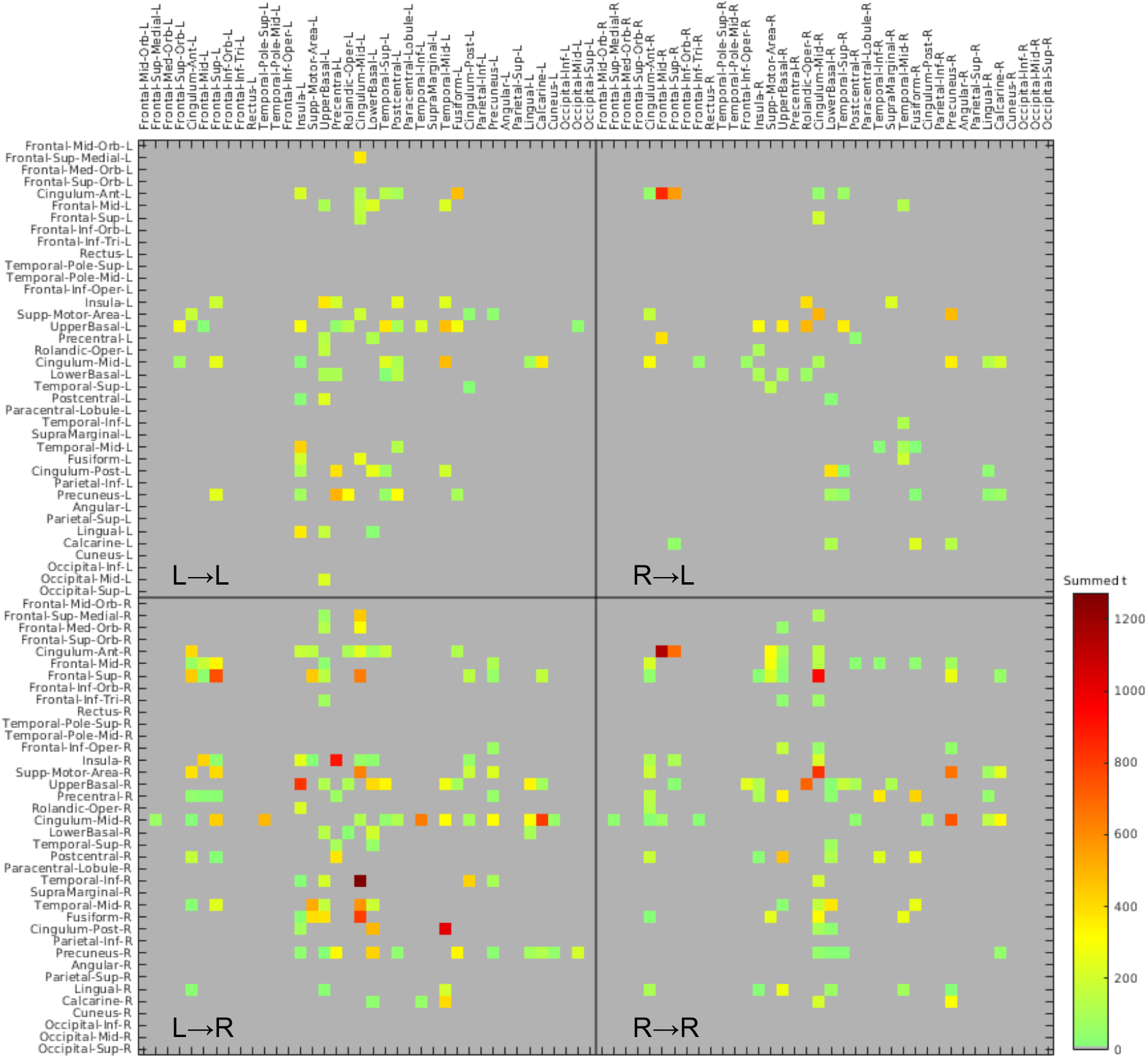
Connectivity matrix of 76 × 76 anatomical regions of interests in schizophrenia patients

### SUPPLEMENTARY MOVIES (Only Notes)

**Supplementary Movie 1–6**

Note: Red arrow indicates high effective connectivity (within 50 ms lag) and blue arrow indicates low connectivity. Connmagnitude and the color indicate amount of inflow. Sphere size and the color indicate amount of total inflow in each node. The bottom panel shows the envelope of the significant edges between 35 and 45 Hz. Abbreviations: RPDC, renormalized partial directed coherence.

